# ElecFeX: A user-friendly toolkit for efficient feature extraction from single-cell electrophysiological recordings

**DOI:** 10.1101/2023.05.27.542584

**Authors:** Xinyue Ma, Loïs S. Miraucourt, Haoyi Qiu, Mengyi Xu, Reza Sharif-Naeini, Anmar Khadra

## Abstract

**Motivation:** Profiling neurons by their electrophysiological phenotype is essential for understanding their roles in information coding within and beyond the nervous systems. Technological development has unleashed our power to record neurons more than ever before, yet the booming size of the dataset poses new challenges for data analysis. Current software tools require users to have either significant programming knowledge or to devote great time and effort, which impedes their prevalence and adoption among experimentalists. To address this problem, here we present ElecFeX, a MATLAB-based graphical user interface designed for a more accessible and efficient analysis of single-cell electrophysiological recordings. ElecFeX has a simple and succinct graphical layout to enable effortless handling of large datasets. This tool includes a set of customizable methods for most common electrophysiological features, and these methods can process multiple files all at once in a reliable and reproducible manner. The output is assembled in a properly formatted file which is exportable for further analysis such as statistical comparison and clustering. By providing such a streamlined and user-friendly open-sourced interface, we hope ElecFeX can benefit broader users for their studies associated with neural activity.

**Summary:** Characterizing neurons by their electrophysiological phenotypes is essential for understanding the neural basis of behavioral and cognitive functions. Recent developments in electrode technologies have enabled the collection of hundreds of neural recordings; that necessitated the development of new toolkits capable of performing feature extraction efficiently. To address this urgent need for a powerful and accessible tool, we present ElecFeX, an open-source MATLAB-based toolbox that (1) has a succinct and intuitive graphical user interface, (2) provides generalized methods for wide-ranging electrophysiological features, (3) processes large-size dataset effortlessly, and (4) yields formatted output for further analysis such as neuronal characterization and classification. We implemented the toolbox on a diverse set of neural recordings and demonstrated its functionality, efficiency, and versatility in capturing features that can well-distinguish neuronal subgroups across brain regions and species. ElecFeX is thus presented as a powerful tool to significantly promote future studies on neuronal electrical activity.

## Introduction

Extracting electrophysiological features is a basic step in a wide range of neuroscience studies. It provides a quantitative description of neural electrical activity which is subsequently used to perform neuronal classification, characterization, and comparison (Cadwell et al., 2016; Gouwens et al., 2019; Zheng et al., 2019). Across species, neurons intermingled in different brain regions are classified based on their molecular, morphological, and transcriptomic profiles and these identified neuronal subgroups also tend to have distinct patterns of electrical activity (Connors and Gutnick, 1990). Indeed, a substantial and influential body of work has relied on neuronal electrical properties to identify neuronal subgroups, especially when the molecular marker is unknown or the transgenic model is not accessible (Nowak et al., 2003; Zaitsev et al., 2012; Ramayya et al., 2014). After identifying neuronal subgroups, the role of each cell type is determined by applying specific perturbations to their neural activity using, for example, pharmacological and molecular manipulations (Fuchs et al., 2007; Zhang et al., 2007). With such perturbations, one can then link their effects bottom-up to behavior and cognitive functions (Bernstein and Boyden, 2011; Sternson and Roth, 2014), as well as top-down to associate them with molecular regulatory pathways that occur subcellularly (Tomaselli and Marbán, 1999; Farjami et al., 2020; McNeill et al., 2021; Ma et al., 2023).

Presently, The way of handling current clamp recordings has been relied on quantitative software. Current software includes AxoGraph X (RRID:SCR_014284), Clampfit (RRID:SCR_011323), and a recent web-based platform NeuroFeatureExtract by EBRAINS (Bologna et al., 2021). These tools have a graphical user interface for intuitive usage and have thus been extensively used among electrophysiologists. However, data analysis using their interfaces is usually conducted in a file-by-file manner, which tends to be laborious, time-consuming, and prone to inconsistency. As the emerging experimental techniques can simultaneously record activities of thousands of neurons (Tsai et al., 2017; Steinmetz et al., 2021), quantifying electrophysiological features has been pushed towards the usage of more powerful and automatic toolkits. Numerous toolboxes are freely accessible and have been developed in varied scientific languages and environments (e.g., MATLAB, Python, R). For instance, pyABF in Python and abf2-package in R have been developed for reading Axon Binary Format (ABF) files from pClamp data acquisition (Harden, 2022; Stanislav and Florian, 2022). Elephant, Pynapple (both in Python), MANTA and PANDORA (both in MATLAB), on the other hand, provide homogeneous and necessary methods for analyzing electrophysiological data and are being constantly maintained and updated by their fervent user communities (Günay et al., 2009; Englitz et al., 2013; Viejo et al., 2022; Denker et al., 2023). However, these command-line based packages target users with significant programming knowledge, which can impede their prevalence among wider user bases.

To address the need of balance between functionality and the accessibility of the toolkit, here we present ElecFeX (<u>elec</u>trophysiological <u>fe</u>ature e<u>x</u>traction toolkit), a MATLAB-based graphical user interface that enables both an intuitive usage and an efficient analysis of multiple electrophysiological recordings. ElecFeX is versatile to process recordings of neurons under different conditions. It provides a sufficient collection of methods to measure most common electrical properties and provides customizable parameters to ensure their flexibility on different signal waveforms. All the settings within the toolkit can be saved to enable the reproduction of the analysis. The recording signals and measured features are simultaneously visualized so that users can tune the parameter settings that best fit their goals of analysis. Once the parameters are confirmed, these measurements can be consistently applied to all recordings in the dataset. The results are then arranged in a table format which is exportable for further analysis. The functionality of ElecFeX in electrophysiological feature extraction was evaluated under a proposed experimental framework commonly adopted in neuroscience research. The versatility of ElecFeX was demonstrated by its implementation on recording datasets collected from different brain regions and species. We showed that ElecFeX successfully captured significant amounts of features to distinguish neuronal subgroups. Together, we provided ElecFeX as a user-friendly and efficient toolkit that advances the analysis of electrophysiological recordings to be efficient, powerful, and reproducible.

## Results

### Electrophysiology-based experimental framework

To address the challenges of quantifying electrophysiological data in neuroscience studies, we developed a toolkit that allows for intuitive and efficient feature extraction from current-clamp recordings. A commonly adopted experimental framework that requires electrophysiological feature extraction is summarized in **Figure 1**. At the initial data acquisition stage (**Figure 1A**), electrophysiological recordings are obtained from numerous neurons under different conditions: from different brain regions, across different species, or under different treatments. To characterize neuronal intrinsic excitability, the electrical responses of neurons were collected in response to current clamp protocols of different waveforms. Given the inherent diversity present in the acquired data, it is imperative for the toolkit to possess a high level of generality. Subsequently, the recorded data are then characterized by a wide range of key features (**Figure 1B**), an indispensable step prior to performing tasks such as data visualization, statistical comparison, or classification (**Figure 1C**). Therefore, an efficient and easy-to-use toolkit like ElecFeX are necessary to serve as an “enzyme” that boosts the feature extraction step, which can be considered as the “rate-limiting” step for the “chain” of investigation on electrophysiology-related questions.

**Figure 1.**
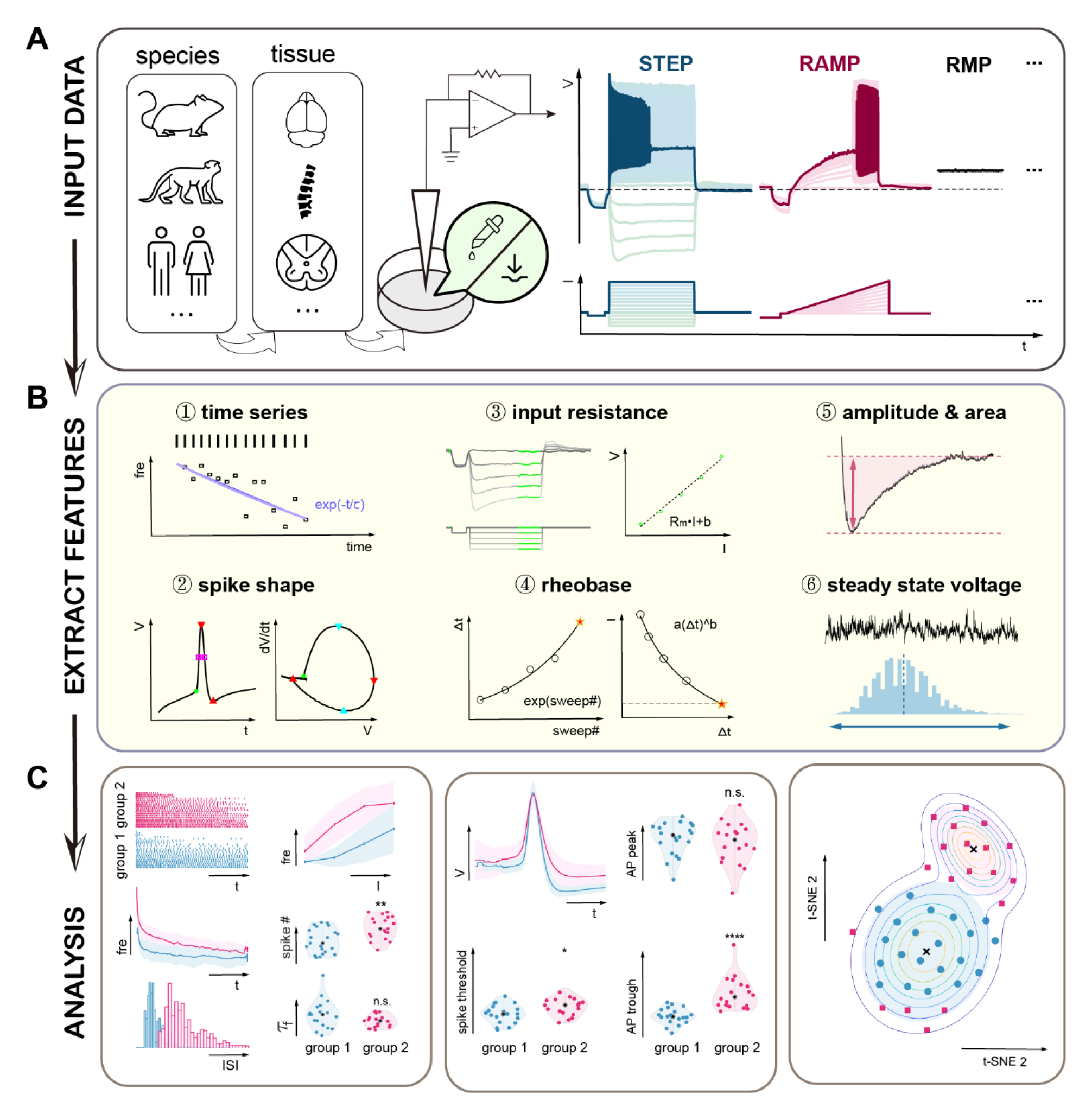
Electrophysiological feature extraction is an indispensable step for a generalized framework of electrophysiology-related studies. **(A)** For the data acquisition, recordings are collected from neurons in different tissues (brain regions or spinal cord) of different species (mouse, marmoset, human, …) under different treatments (chemical, mechanical, …). To characterize the electrical properties, neurons are stimulated by injected currents of different waveforms, such as step, ramp, or I=0 (RMP). **(B)** A collection of electrophysiological features that can be extracted from current-clamp recordings. **(C)** The analysis of the electrophysiological features further provides information between the known (statistical comparison) or unknown (clustering) groups.

### Graphical user interface and pipeline

The graphical user interface (GUI) of ElecFeX (**Figure 2**) is designed to be user-friendly with texts that guide the user through the complete feature extraction workflow. The main GUI window encompasses all the essential elements required for the analysis process, and there is another callable window for advanced settings related to measuring spike properties. The main window of GUI is organized in accordance with the analysis procedures and is divided into six sections: (1) load file, (2) data info, (3) feature extraction, (4) visualization, (5) batch analysis setting, and (6) export results. Detailed instructions regarding installation and implementation of the toolkit can be found on the corresponding GitHub page https://github.com/XinyueMa-neuro/ElecFeX.

**Figure 2.**
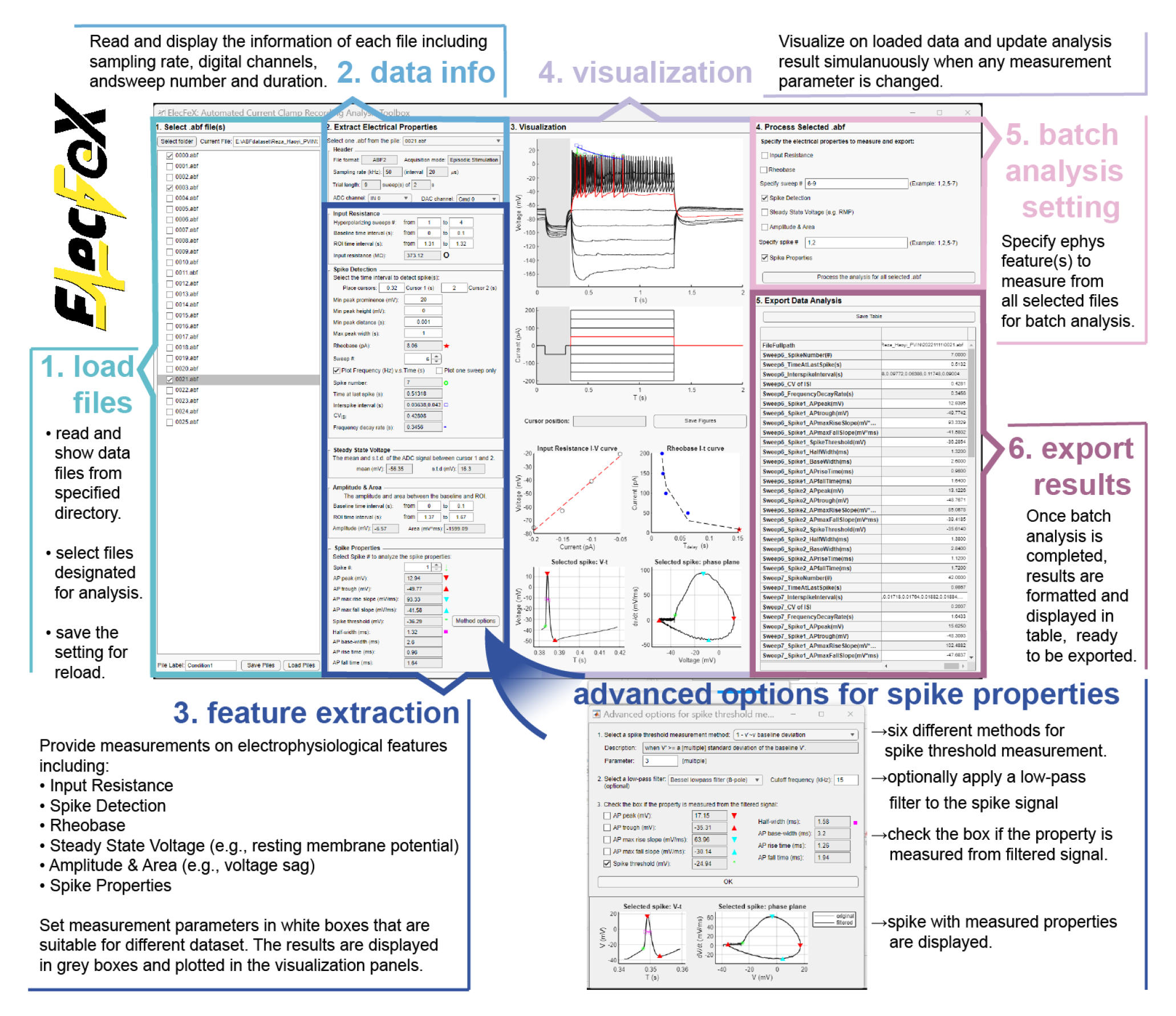
A layout description of ElecFeX’s graphical user interface (GUI). The GUI of ElexFeX contains six different panels classified based on the usage steps: (1) load files from the dataset, (2) display each data information, (3) measure electrophysiological features, (4) visualize data and extracted features, (5) set for multiple file processing (batch analysis), and (6) export results.

The user begins by inputting the directory where the dataset is located. All data files in Axon™ Binary File format (ABF) within the specified directory are loaded and displayed in a hierarchical structure mirroring their file locations. Each file is accompanied with a selectable checkbox, which allows the user to designate files as a pile for later batch analysis; the file selection settings can be saved for future usage.

The subsequent sections serve to read, visualize, analyze each file and determine the optimal parameter settings for electrophysiological feature extraction. The “data info” section displays the file information including file format, acquisition mode, sampling rate, sweep number and duration, and signal channels, which corresponds to the protocol setting in pClamp data acquisition software. The drop-down menu of analog-to-digital converter (ADC) and digital-to-analog converter (DAC) channels enables the selection of the desired channel from which signals will be analyzed as response and stimulus signals.

ElecFeX provides a collection of methods to extract commonly studied electrophysiological features. These methods are equipped with customizable parameters (indicated as white box texts in Figure 3), allowing for flexibility in processing signals with various waveforms. Details of the quantification methods are described in the following section “Electrophysiological feature extraction methods” and “Electrophysiological feature analysis” of “STAR methods” section.

**Figure 3.**
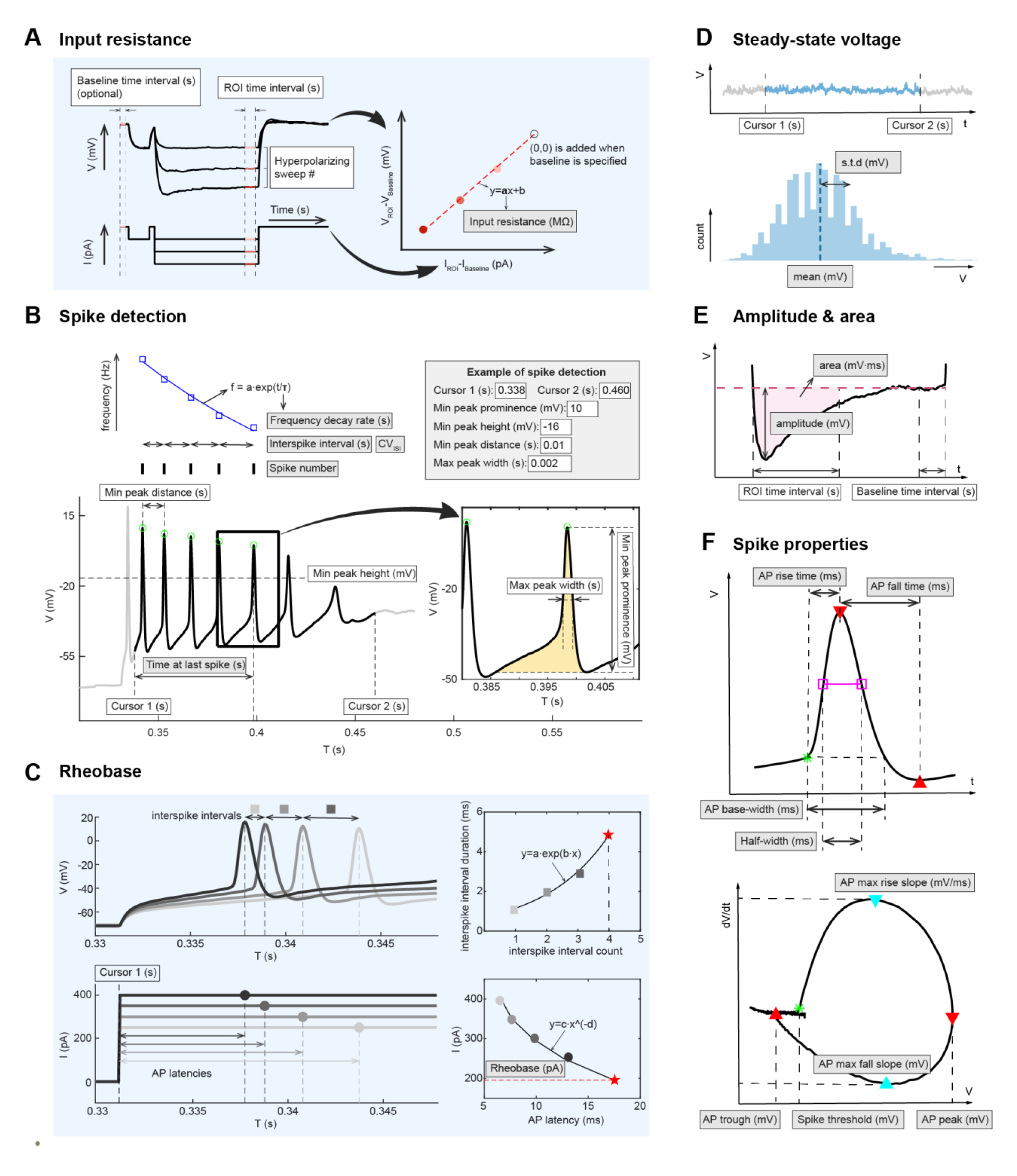
Six categories of electrophysiological features methods with parameter settings provided by ElecFeX. **(A)** An example of input resistance measurement. The users are required to specify parameters (white boxes) including the sweeps, time intervals of baseline and region of interest (ROI). The sampled datapoints (ROI-baseline) of voltage against that of current is fitted by a linear function, where the slope of the curve is displayed in the grey box named “input resistance (MΩ)”. **(B)** All the spikes that (1) is between “Cursor 1” and “Cursor 2” and (2) meet the customized criteria (all the texts in white boxes) are detected. The top-right panel is an example parameter setting for spike detection and the detected spikes are marked by green circles. Several quantifications (texts in grey boxes) on the time series of the spike trains are conducted, such as the “spike number” and “interspike intervals”. The instantaneous firing frequencies (purple squares) are measured and fitted to a single exponential decay function (purple curve), where the time constant refers to the “frequency decay rate”. **(C)** The description of rheobase measurement. For each sweep, the time interval from the first spike (if any) to the initiation of test stimuli (specified by “Cursor 1”), called action potential (AP) latency is measured. The interspike intervals of these first spikes (grey filled squares) are fitted to a single exponential function (top-right panel), from which the expected AP latency of first spike (red pentagram) triggered by one-step size smaller current is estimated. The current values against AP latencies (grey filled circles) are then fitted to a power function (bottom-right panel) and the rheobase is estimated by expected AP latency. **(D)** The measurements of “steady state voltage”. The average value (“mean (mV)”) and standard deviation (“s.t.d. (mV)”) of voltage between “Cursor 1” and “Cursor 2” are measured. **(E)** The amplitude and area are measured as the difference between the region of interest (ROI) and the baseline voltage. **(F)** Multiple spike properties measured from the spike shape (top panel) and the AP cycle (bottom panel).

The extracted features are simultaneously visualized in the "visualization" section, facilitating convenient modification of parameter settings until the desired analysis results are achieved. Once the optimal method parameter settings have been determined, the user can proceed to the "batch analysis setting" section to specify the desired sweeps or spikes on which the methods will be applied. Once all settings have been finalized, the batch analysis is ready to be executed, during which all selected files are consistently processed using the specified methods. The results are then formatted and presented in a table format and can be exported as an Excel spreadsheet for further analysis purposes.

### Electrophysiological feature extraction methods

To minimize the usage complexity but maximize the attainability of features from diverse signal waveforms, we generalized the electrophysiological feature extraction methods into six categories and provided customizable parameters for each method (**Figure 3**). For instance, the “input resistance” subsection measures the slope of a linear fit between the sampled DAC and ADC signal (**Figure 3A**); “steady state voltage” subsection measures the average and standard deviation of sampled ADC signal (**Figure 3D**); “amplitude & area” subsection quantifies the strength of any transient change in ADC signal (**Figure 3E**). ElecFeX ensures a tailored fit of each method to the specific data being processed.

We also incorporated a novel approach for estimating the rheobase from step-current recordings. The rheobase is determined by identifying the injection currents at which the first spike exhibits a time latency exponentially greater than the larger injection currents (**Figure 3B**). This method guarantees a consistent and standardized estimation of the rheobase, regardless of the specific step-current protocol employed.

Another crucial aspect of feature extraction involves spike detection and isolation. Spike detection is performed by “findpeaks()” function from MATLAB Signal Processing Toolbox. Leveraging the detected spike trains, ElecFeX provides a comprehensive characterization of the time series, including spike number, discharge duration, interspike intervals, and frequency decay rate (**Figure 3B**). Each spike can be isolated from the spike train and described through a set of properties derived from the temporal sequence and phase plane of the spike signal (**Figure 3F**). In the advanced setting for spike properties, six different ways of measuring spike thresholds were adopted from previous work (Sekerli et al., 2004), such as the voltage at which dv/dt reaches an ad hoc value. This integration allows users to select the method that aligns best with their familiarity or preference. A suite of low-pass filters is also provided to refine the analysis outcomes, if needed, thereby accommodating specific requirements and enhancing the quality of the results.

### Region-specific variance in neuronal electrical properties is quantitatively captured by ElecFeX

To evaluate the functionality and efficiency of ElecFeX, we conducted a series of implementations using different electrophysiological datasets. The first dataset contains whole-cell patch-clamp recordings of parvalbumin-expressing neurons (PVn) from mice spinal dorsal horn (SDH) and hippocampus (HPC). The PVns are identified as fluorescent positive neurons from PVcre:tdTom mice. Neural recordings were collected in response to three standardized current-clamp protocols (step, ramp, and I=0) and only those failed to meet predefined quality control criteria were excluded for further analysis (**Figure 4A**).

**Figure 4.**
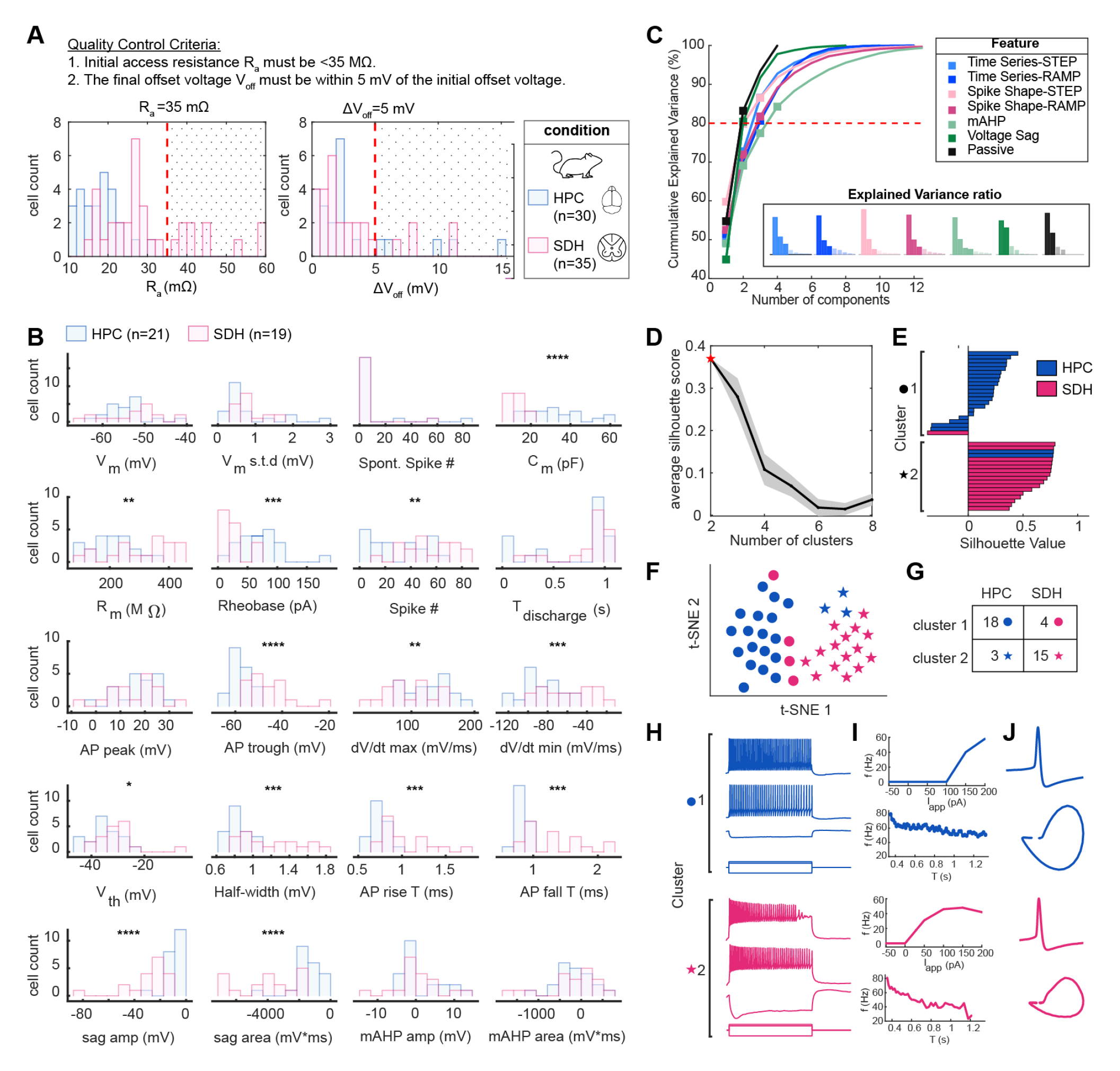
ElecFeX extracts electrical features that can distinguish parvalbumin-expressing neurons (PVns) from different regions of the nervous systems. **(A)** The recordings of PVns from the hippocampus (HPC) and the spinal dorsal horn (SDH) underwent quality control given the access resistance and the offset voltage before and after the stimulus and those failed to meet the criteria (black dotted areas in the histogram) were excluded from the dataset. **(B)** Histogram distributions for electrophysiological features extracted by ElecFeX. The distributions of the 20 electrophysiological features of PVns from HPC (blue) and SDH (pink) were displayed and compared by unpaired parametric t-test. **(C)** Dimensionality reduction of electrophysiological features in seven feature categories. Principal component analysis was performed on the data subset that belongs to each feature category. The preceding components that cumulatively explain 80% of the variance (red dashed line) were collected for further clustering. Insets are bar plots of explained variance ratio of identified components, of which the solid color refers to the collected components and the half-transparent color refers to rejected components. **(D)** To determine the optimal number of clusters for k-means clustering, the average silhouette score (black line; the grey area is mean±SEM) was calculated for k-means clustering analyses using between 2 and 8 clusters. The highest silhouette score was achieved using two clusters (red pentagram). **(E)** The silhouette value for each neuron is shown in their corresponding cluster determined by k-means clustering. Low or negative silhouette values indicate points that fit poorly within its cluster. **(F)** t-SNE plots for the clustering result. Different colors refer to the conditions and different marker shapes refer to the cluster. **(G)** Quantification of the number of PVns from different tissues that are clustered into different subgroups **(H)** Representative traces of neurons from cluster 1 and 2. **(I)** The f-I curves (top) and the instantaneous frequencies (bottom). **(J)** The shape and AP cycles of the first spike triggered by a step current stimulus.

The final dataset comprised a total of 21 hippocampal PVns and 19 spinal dorsal horn PVns. We implemented ElecFeX to fully extract electrophysiological features from this dataset and generated a feature matrix (**Table 1**). The distributions of 20 representative electrophysiological features from ElecFeX were presented and compared between HPC and SDH PVns (**Figure 4B**). The cell body size of HPC PVns were observed to be larger than SDH PVns (data not shown); such morphological differentiation was consistently reflected in the disparities observed in electrical properties such as whole cell capacitance, input resistance, and rheobase. Moreover, significant differences were observed in spike shape between HPC and SDH PVns, suggesting their specialized roles in information coding within distinct neural circuits across different regions of the nervous system.

**Table 1.** Electrophysiological feature data sets for PVns

| Name | Description | Principle components used |
| --- | --- | --- |
| Time Series-STEP | Spike number, discharge duration, $CV_{ISI}$, and frequency decay rate. From all the depolarizing step sweeps. | 3 |
| Time Series-RAMP | Spike number, discharge duration, $CV_{ISI}$, and frequency decay rate. From all the ramp sweeps. | 3 |
| Spike Shape-Step | AP peak, AP trough, spike threshold, AP max rise slope, AP max fall slope, half-width, base-width, AP rise time, and AP fall time. From first 25 spikes (if any) of all the depolarizing step sweeps. | 3 |
| Spike Shape-Ramp | AP peak, AP trough, spike threshold, AP max rise slope, AP max fall slope, half-width, base-width, AP rise time, and AP fall time. From first five spikes (if any) of all the ramp sweeps. | 3 |
| mAHP | mAHP from all the depolarizing sweeps (both step and ramp) | 4 |
| Voltage sag | Voltage sag from all the hyperpolarizing step sweeps. | 2 |
| Passive | Whole cell capacitance, input resistance, rheobase, resting membrane potential, and standard deviation of resting membrane potential. | 2 |

We further explored the significance of ElecFeX’s output; this was demonstrated by performing neuronal subgroup clustering based on their electrophysiological feature-matrix extracted from ElecFeX. Given the complexity and interrelationships of several features within the matrix, we first categorized all the features into seven groups (**Table 1**) and specifically reduced their dimensionality by principal component analysis (PCA) (**Figure 4C**). For instance, “Time series-STEP” composes all the descriptive properties of the time series of spike trains that were collected from the voltage response of each depolarizing step and ramp currents for each cell. Following PCA, we collected the principal components which explained variance exceeded a defined threshold, resulting in the collection of 20 PCA features. These PCA features were then used for K-means clustering analysis, with the optimal number of clusters determined based on the maximum average silhouette value (**Figure 4D**). The clustering results were visualized in the silhouette plots for each cell (**Figure 4F**) and by projecting PCA features onto two dimensions using the t-distributed stochastic neighbor embedding (t-SNE) method (van der Maaten and Hinton, 2008) (**Figure 4F-G**). Notably, the projection demonstrated that clustering based on ElecFeX-derived electrophysiological features separated the HPC PVns and SDH PVns reasonably well, especially when the difference between representative traces of two clusters are not visually pronounced (**Figure 4H-J**). These results demonstrate the efficiency of ElecFeX in capturing electrophysiological features that enable the classification of neuronal subgroups.

### ElecFeX captures electrophysiological features that consistently distinguish neuronal subgroups across species

The versatility of ElecFeX was further examined by a different set of recordings obtained from mice and marmoset dorsal root ganglions (DRGs). DRGs are the first-order neurons in the somatosensory afferent pathway and their electrical properties are drastically different than PVns. More importantly, DRGs have been found as a mixture of neuronal subgroups exhibiting a wide range of electrophysiological properties (Zheng et al., 2019). Therefore, we investigated if ElecFeX is capable of processing diverse datasets and more specifically, generating sufficient features to identify DRG subgroups across different species.

After the quality control according to the access resistance of the recordings, 40 mice DRGs and 25 marmoset DRGs were considered to be processed by ElecFeX and the distributions of 21 representative feature extracted were illustrated in **Figure 5A**. The clustering procedures were similar in the previous section (**Figure 5B-F**). Given the prior knowledge of DRGs, we aimed to maximize the cluster numbers while maintaining the cluster quality. This was determined by identifying the point at which the silhouette value exhibited a steep decrease (**Figure 5C**). Four clusters were identified from the dataset, with each cluster consisting of a mixture of both mice and marmoset recordings (**Figure 5E-F**). The identity of the resulted clusters can be further predicted based on the electrical profiles of DRG subgroups identified in previous work (Zheng et al., 2019) (**Figure 5G-J**). For instance, the clusters exhibiting more active firing behaviors (orange and green) closely resembled the profiles associated with Aδ- and C-fibers and turned out to consistently have lower whole cell capacitance (**Figure 5G-H**). Taken together, these results not only support the functionality of ElecFeX in handling diverse sets of electrophysiological data, but also serve as an example as to how to incorporate ElecFeX into the work of characterizing and classifying neuronal subgroups. The methods for neuronal subgroup clustering are currently provided as customized code in https://github.com/XinyueMa-neuro/ElecFeX-MaEtAl2023 and will be included in the future release or extension of ElecFeX.

**Figure 5.**
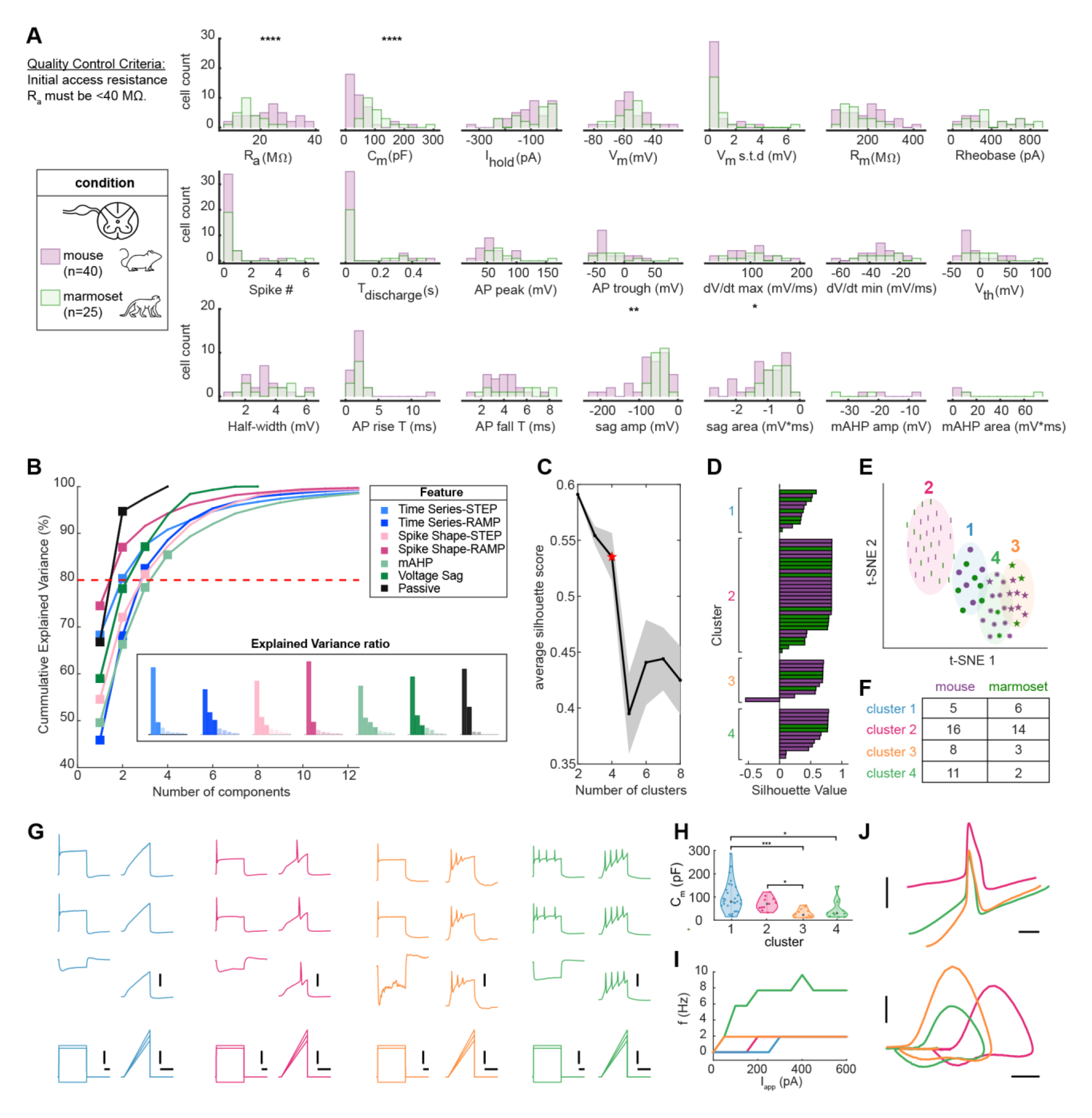
The identification of neuronal subgroups of DRGs from different species based on ElecFeX-generated electrophysiological properties. **(A)** Histogram distributions for electrophysiological features extracted by ElecFeX. The distributions of the 21 electrophysiological features of PVns from mice (purple) and marmosets (green) are displayed and compared by unpaired parametric t-test. The recordings that failed to meet the quality control criterion were excluded from the dataset. **(B)** Dimensionality reduction of seven subsets of electrophysiological features by principal component analysis. The preceding components that cumulatively explain 80% of the variance (red dashed line) were collected for further clustering. Bar plots in the inset are the explained variance ratio of identified components. For each bar, the solid and half-transparent colors refer to the collected and rejected components. **(C)** The average silhouette score (black line; the grey area is mean±SEM) was calculated for k-means clustering analyses using between 2 and 8 clusters. The cluster number n=4 (red pentagram), at which silhouette score starts to decrease sharply were used. **(D)** The silhouette value for each neuron is shown in their corresponding cluster determined by k-means clustering. **(E)** t-SNE plots for the clustering result. Different color refers to conditions and different marker shapes refers to the cluster. **(F)** Quantification on the number of DRGs from different species that are clustered into different subgroups **(G)** Representative traces of DRGs from cluster 1-4. Scale bars: top vertical bars, 60 mV; middle vertical bars, 500 pA; bottom horizontal bars, 0.1 s. **(H)** The comparison of whole cell capacitance between different clusters. **(I)** The f-I curves. **(J)** The shape and AP cycles of the first spike triggered by ramp current stimulus. Scale bars: top vertical bar, 50 mV; top horizontal bar, 0.1 s; bottom vertical bar, 50 mV/ms; bottom horizontal bars, 30 mV.

## Discussion

Here, we introduced ElecFeX, a user-friendly and efficient open-source toolbox for electrophysiological feature extraction. Quantifying electrophysiological features are imperative in explaining many experimental observations to studies of different levels, from molecular regulation pathways within neurons to the neural basis of higher brain functions. To facilitate this procedure, the toolkit includes a graphical user interface to facilitate intuitive usage and enables quality control of analysis results. Users can customize method settings to accommodate diverse datasets, facilitating flexible and reliable processing of multiple files. The file and parameter settings can also be saved and retrieved, ensuring reproducibility of analysis results.

At the core of ElecFeX lies in its ability to offer robust and versatile analysis of electrophysiological data. ElecFeX minimizes usage complexity while maximizing accessibility to a wide array of features and accommodating various signal waveforms. To achieve this, the electrophysiological feature extraction methods are generalized into six different categories (**Figure 3**), with each method accompanied with parameters to be set by the users. The functionality, efficiency, and versatility of ElecFeX were demonstrated through its successful application to two different sets of electrophysiological recordings in whole-cell configuration. These results exemplify the significant value of ElecFeX in capturing electrical features that differentiate neuronal subgroups, showcasing its immense potential in advancing future electrophysiology-related investigations.

In conclusion, ElecFeX stands as a user-friendly, powerful open-source toolkit for feature extraction, adaptable to diverse electrophysiological data. The toolkit ElecFeX presented in this paper offers significant advantages for investigating critical questions regarding electrophysiology-related studies. Its main advantages over several existing toolboxes are the ease of use and simplicity, the generalization and comprehensiveness of measurement methods, the versatility to adapt diverse datasets, and the efficiency and reproducibility in batch analysis. The toolkit has already been shared with and beyond our labs, garnering feedback, and suggestions for improvement. Ultimately, our goal is to ensure that our toolkit indeed promotes the analysis experience for the community. We strongly encourage more engagement and collaboration from the neuroscience community to foster the development of ElecFeX and serve the evolving needs of the neuroscience field.

### Limitations of Study

While the presented tool offers great advantages for investigating electrophysiological data, there are limitations that can be addressed and updated in future releases. As discussed earlier, ElecFeX is currently specific for the data format of .abf (Axon Binary File), which is the original file format from pClamp data acquisition software. To ensure extensive usage, we are aiming to extend the data formats of the files that the user can upload and process (e.g., data format from other data acquisition software like Igor, and the neurodata without boundary (nwb); all electrophysiological data formats accepted by the BluePyEfe and the Neo package; Garcia et al., 2014). Another potential barrier for prevalent usage is that ElecFeX is currently only available in MATLAB. We have addressed the concern of MATLAB dependency by providing a standalone desktop application format of ElecFeX so that users can access ElecFeX without the need to have MATLAB installed in their operating system.

ElecFeX expands the accessibility of data analysis by incorporating customized code with GUI, but it can also lead to limitations inherent to GUI. First, despite providing as many parameter settings as possible for customizable measurements, GUI is still not as flexible as command-line tools or scripts. While they can be very user-friendly and easy to use, they may not offer the same level of control and customization that more advanced users require. This could be a disadvantage for researchers who want to develop more complex analyses or custom scripts. Second, GUIs can sometimes be slower than command-line tools or scripts, particularly when working with large datasets or complex analyses. This is because GUIs often rely on a graphical rendering engine to display visualizations, which can be resource intensive. This could be a disadvantage for researchers who need to process large amounts of data quickly or who want to perform real-time analyses. We managed to accelerate the speed for multiple file processing by activating MATLAB-based parallel processing and future work will test the speed of ElecFeX in processing large-size datasets, e.g., electrophysiological recordings from Allen Institute. Future work will evaluate the speed of our toolkit compared with other available software packages.

## Acknowledgments

This work was supported by a project grant from the Canadian Institutes of Health Research to R.S.N. (CIHR PJT-162404) and the Natural Sciences and Engineering Council of Canada (https://www.nserc-crsng.gc.ca/index_eng.asp) discovery grant to A.K. and the McGill Initiative in Computational Medicine (https://www.mcgill.ca/micm/) team grant to A.K. and R.S.N. X.M. was supported by the Globalink Graduate Fellowship. We thank the lab members in R.S.N.’s lab for testing the toolkit and providing valuable feedback.

## Author contributions

Conceptualization, X.M. and L.M.; Methodology, X.M., L.M., H.Q., and M.X.; Software, X.M.; Formal Analysis, X.M.; Investigation, X.M., L.M., H.Q., and M.X.; Writing – Original Draft, X.M.; Writing – Review & Editing, X.M., L.M., R.S.N., and A.K.; Supervision, R.S.N., and A.K.; Funding Acquisition, R.S.N., and A.K.

## Declaration of Interests

The authors declare no competing financial interests.

### STAR Methods

### Key resources table

STAR Methods Key resources table
| REAGENT or RESOURCE | SOURCE | IDENTIFIER |
| --- | --- | --- |
| **Deposited data** |  |  |
| An example dataset of recordings from mice parvalbumin-expressing neurons | This paper | GitHub: https://github.com/XinyueMa-neuro/ElecFeX-MaEtAl2023 |
| Code for the toolkit ElecFeX | This paper | GitHub: https://github.com/XinyueMa-neuro/ElecFeX |
| Code for analysis reproduction | This paper | GitHub: https://github.com/XinyueMa-neuro/ElecFeX-MaEtAl2023 |
| **Experimental models: Organisms/strains** |  |  |
| Mouse: B6.129P2-*Pvalb*<sup>tm1(cre)Arbr/J</sup> | The Jackson Laboratory | https://www.jax.org/strain/017320 |
| Mouse: B6.Cg-*Gt(ROSA)26Sor*<sup>tm14(CAG-tdTomato)Hze/J</sup> | The Jackson Laboratory | https://www.jax.org/strain/007914 |
| Mouse: PVcre;tdTom | Haoyi et al. | N/A |
| Mouse: C57/BJ mouse |  | https://www.jax.org/strain/005304 |
| Marmoset | This paper | N/A |
| **Software and algorithms** |  |  |
| ElecFeX | This paper | GitHub, |
| MATLAB R2022a | Mathworks | RRID:SCR_001622,<br>http://www.mathworks.com/products/matlab/ |
| MultiClamp 700B Microelectrode Amplifier | Molecular Devices | RRID:SCR_018455 |
| Molecular Devices DigiData 1440 digitizer | Molecular Devices | RRID:SCR_021038 |
| pClamp | Molecular Devices | RRID:SCR_011323 |

### Resource availability

#### Lead contact

Further information and requests for resources and reagents should be directed to and will be fulfilled by the lead contact, Xinyue Ma.

#### Material availability

This study did not generate new unique reagents.

#### Data and code availability

- An example dataset of recordings from mice parvalbumin-expressing neurons has been deposited at GitHub and is publicly available as of the date of publication. DOIs are listed in the key resources table. All electrophysiological data reported in this paper will be shared by the lead contact upon request.
- The toolkit and all original code to reproduce the figures has been deposited at GitHub and is publicly available as of the date of publication. DOIs are listed in the key resources table.
- Any additional information required to reanalyze the data reported in this paper is available from the lead contact upon request.

### Experimental model and subject details

#### Mice

All mice were kept under pathogen-free conditions, and cages were maintained in ventilated racks in temperature (20–21 ℃) and humidity (55%) controlled rooms on a 12 h light/dark cycle, with food and water provided *ad libitum*. All experimental procedures were approved by the Animal Care and Use Committee at McGill University, in accordance with the regulations of the Canadian Council on Animal Care. The PV^cre^;tdTom mice were generated by crossing commercially available PV^cre^ mice (JAX, stock #017320, https://www.jax.org/strain/017320) with Ai14 tdTomato reporter (JAX, stock#007914, https://www.jax.org/strain/007914). The hippocampal PVn recordings were from 8 male mice and the spinal dorsal horn PVn recordings were from 5 female and 8 male mice. The DRGs were from 6 female and 2 male mice. Sex was not considered as a variable in this study.

#### Marmosets

Three (1 male and 2 females) common marmosets (*Callithrix jacchus*; McGill University breeding colony) weighing 300-450 g and aged between 2 and 6 years old during the experiments were used. Marmosets were housed in groups of 2 under conditions of controlled temperature (24 ± 1 °C), humidity (50 ± 5%) and a 12 h light/dark cycle (07:15 lights on). They had unlimited access to water; food (Mazuri® Marmoset Jelly, boiled eggs, boiled pasta, nuts, legumes) and fresh fruits were served twice daily. Their home cages were enriched with primate toys and perches. Animals were cared for in accordance with a protocol approved by McGill University and Montreal Neurological Institute-Hospital (The Neuro) Animal Care Committees (Animal Use Protocol 2017–7922), both in accordance with regulations defined by the Canadian Council on Animal Care.

### Method details

#### Slice Preparation

Adult mice (6-8 weeks old) were anesthetized with 5% isofluorane and perfused transcardially with ice-cold oxygenated (95% O2, 5% CO2) N-Methyl-D-glutamine-based artificial cerebrospinal fluid (NMDG-ACSF) solution (93 NMDG, 93 HCl 12M, 2.5 KCl, 1.25 NaH2PO4, 30 NaHCO3, 20 HEPES, 25 glucose, 2 thiourea, 5 Na-L-ascorbate, 3 Na-pyruvate, 10 N-acetyl-L cysteine, 0.5 CaCl2/2H2O, and 10 MgSO4/7H2O, in mM; 300±10 mOsm; pH 7.3–7.4). The lumbar spinal segment was rapidly removed and immersed in ice-cold NMDG-ACSF. The ventral roots were removed and 300 µm transverse slices were cut with a vibratome (Leica VT1000S). Slices were transferred to a submerged chamber containing HEPES-based recovery ACSF (92 NaCl, 2.5 KCl, 1.25 NaH2PO4, 30 NaHCO3, 20 HEPES, 25 Glucose, 2 Thiourea, 5 Na-ascorbate, 3 Na-pyruvate, 2 CaCl2, 2 MgSO4, in mM; 300±10 mOsm, pH 7.4) for at least 10 minutes at 34- 36 ℃, equilibrated with 95% O2 and 5% CO2. At the end of the slicing period, the HEPES-based recovery ACSF was let to equilibrate at room temperature for at least one hour.

#### Marmoset DRGs dissection and culture

Marmoset DRGs were from the lumbar vertebra. The connective and fat tissue around DRGs were removed in ice-cold NMDG-aCSF. Ganglia were enzymatically digested at 37°C for 1h with a Dispase II / type IV Collagenase mixture dissolved in DMEM/F12. Ganglia were mechanically dissociated by gentle trituration then filtered through a 100 μm cell strainer and seeded on to culture dishes. Cells were maintained at 37°C with 5% CO2 in Neurobasal A media supplemented with 5% Fetal bovine serum, 2% Neuroculture SM1 Neuronal Supplement, 1x Penicillin-Streptomycin, 1x GlutaMax and 25 ng/ml NGF. Half of the culture media was replaced with fresh media every 3 days.

#### Mouse DRGs dissection and culture

Mouse DRGs were collected from the lumbar vertebra (C57/BJ mouse, JAX, stock #005304, https://www.jax.org/strain/005304). Ganglia were enzymatically digested at 37°C for 0.5h with a Dispase II / type IV Collagenase mixture dissolved in DMEM/F12. Ganglia were mechanically dissociated by gentle trituration then seeded on to culture dishes. Cells were maintained at 37°C with 5% CO2 in DMEM/F12 supplemented with 10% Fetal bovine serum, 1x Penicillin-Streptomycin and 25 ng/ml NGF. Half of the culture media was replaced with fresh media every 2 days.

#### Electrophysiology

For slice recordings, slices were transferred individually to a recording chamber. The slices and the primary cultured cells were kept at room temperature and were continuously superfused with oxygenated ACSF (126 NaCl, 26 NaHCO3, 2.5 KCl, 1.25 NaH2PO4, 2 CaCl2, 2 MgCl2, and 10 glucose, in mM; bubbled with 95% O2 and 5% CO2; pH 7.3; osmolarity, 300±10 mOsm measured, 2 ml/min). Patch pipettes were pulled from borosilicate glass capillaries (Harvard Apparatus) with a P-97 puller (Sutter Instruments). They were filled with a solution containing 135 K-gluconate, 6 NaCl, 0.1 EGTA, 10 HEPES, 2 MgCl2, 2 Mg-ATP, and 0.8 Na-GTP (in mM; pH 7.4, adjusted with KOH; osmolarity, 300±10 mOsm, adjusted with sucrose) and had final tip resistances of 6–8 MΩ for whole cell recording. Holding potentials were applied to correct a liquid junction potential of −15.9 mV and it was not adjusted for all the voltage properties presented in this study. Neurons were viewed by an upright microscope (Olympus) with a 40X water-immersion objective, infrared differential interference contrast (IR-DIC) and fluorescence. For slice recordings, cells expressing the tdTomato were identified as PV-neurons. Data was acquired with pClamp 10 suite software (Molecular Devices) using MultiClamp 700B patch-clamp amplifier and Digidata 1440A (Molecular Devices). Recordings were low pass filtered on-line at 1 kHz, digitized at 20 kHz.

For the slice recordings, three standardized current-clamp protocols were applied per cell recording: (1) a gap-free 90-s long recording with no injection current (I=0) was used to measure the resting membrane potential (mV), (2) a 1-s long step current injection from −200 pA to 200 pA with a step size of 50 pA, and (3) a 1-s long ramp current injection of varied slopes from 50 pA/s to 400 pA/s. For primary cultured cells, the three standardized current-clamp protocols were (1) a gap-free 90-s long recording with no injection current (I=0) was used to measure the resting membrane potential (mV), (2) a 0.5-s long step current injection from –0.2 nA to 1.2 nA with a step size of 0.1 nA, and (3) a 200-ms long ramp current injection of varied slopes from 0.5 nA/s to 9 nA/s.

#### Electrophysiological feature analysis

All the passive and active membrane properties (**Table 1&2**) were collected from ElecFeX and their definitions are provided as follows. Resting membrane potential was measured as the average of voltage between 35 s and 45 s period from a 90-s recording with no injection current. Input resistance was obtained from the slope of the I-V curve in response to hyperpolarized injection currents. Whole-cell capacitance was determined by the amplifier’s automated whole-cell compensation function. Rheobase was defined as the minimal injected step current value needed to evoke a single action potential. We plotted the latency of the first spike against the injected step current value, fit the datapoint to power function *I*_*app*_ = *t*^−*a*^ + *b* where the rheobase was measured as the bottom asymptote y=b that cross Y-axis. The spike threshold, measured as the point at which 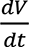 deviated from the mean of the baseline by 3 standard deviations. AP peak is the maximum potential and AP trough is the minimum potential in the interval between two consecutive AP peaks. Half-width is the AP duration at the membrane voltage halfway between the spike threshold and AP peak. AP base-width is the AP duration at the membrane voltage equals to spike threshold. AP rise and fall times were calculated between the voltages from spike threshold to AP peak. The area of medium after-hyperpolarization potential (mAHP) was measured as the area between the resting voltage and the voltage within 200 ms after the test stimulus. The amplitude of mAHP is the difference between the resting voltage and the minimal voltage within 200 ms after the test stimulus. Firing frequencies were measured as the reciprocal of the inter-spike intervals. Its time course was well fitted by a single exponential function *f*(*t*) = *f*_*ss*_ + (*f*_0_ − *f*_*ss*_). exp 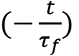, where *τ*_*f*_ denotes the spike frequency decay rate.

#### Electrophysiological classification

Feature subsets were built by accumulating the feature vectors in each category (for example, passive membrane properties, properties of time series and spike shapes from step and ramp current recordings, medium after-hyperpolarization, and voltage sag; see **Table 1&2**). The missing values in the matrix were assigned as an arbitrary value. Principal component analysis (PCA) was separately performed on each feature subset. The preceding principal components with cumulative explained variance exceeding 80% were kept (typically one to four components from a given subset). The components were then z-scored to standardize the scale and combined to form a reduced dimension feature matrix. The matrix was then used to perform K-means clustering analysis. We iteratively performed K-means clustering 10 times for the cluster number between 2 and 8. The optimal number of clusters was determined when the average silhouette value is maximal or starts to drop steeply. The feature matrix used in the clustering analysis was also visualized with a 2D projection using the t-SNE technique.

**Table 2.** Electrophysiological feature data sets for DRGs.

| Name | Description | Principle components used |
| --- | --- | --- |
| Time Series-STEP | Spike number, discharge duration, $CV_{ISI}$, and frequency decay rate. From all the depolarizing step sweeps. | 2 |
| Time Series-RAMP | Spike number, discharge duration, $CV_{ISI}$, and frequency decay rate. From all the ramp sweeps. | 3 |
| Spike Shape-Step | AP peak, AP trough, spike threshold, AP max rise slope, AP max fall slope, half-width, base-width, AP rise time, and AP fall time. From first three spikes (if any) of all the depolarizing step sweeps. | 3 |
| Spike Shape-Ramp | AP peak, AP trough, spike threshold, AP max rise slope, AP max fall slope, half-width, base-width, AP rise time, and AP fall time. From first five spikes (if any) of all the ramp sweeps. | 2 |
| mAHP | mAHP from all the depolarizing sweeps (both step and ramp) | 4 |
| Voltage sag | Voltage sag from all the hyperpolarizing step sweeps. | 3 |
| Passive | Whole cell capacitance, input resistance, rheobase, resting membrane potential, and standard deviation of resting membrane potential. | 2 |

#### Statistical Analysis

Data were expressed as mean±SEM, with n being the number of cells. When comparing between two groups of cells, a 2-tailed unpaired parametric t-test was applied. Statistical analysis was performed in MATLAB R2022a. P≤0.05 (*) was considered to indicate a statistically significant difference, with p<0.01 (**), p<0.001 (***) and p<0.0001 (****) also be noted.

### Quantification and statistical analysis

Details of the quantification and statistical methods used in this study, such as electrophysiological feature analysis and statistical comparison, are described in detail in the respective subsections of the “method details” section within the STAR methods and in the appropriate figure legends.

